# Drug repurposing through joint learning on knowledge graphs and literature

**DOI:** 10.1101/385617

**Authors:** Mona Alshahrani, Robert Hoehndorf

**Affiliations:** Computer, Electrical and Mathematical Sciences and Engineering Division, Computational Bioscience Research Center, King Abdullah University of Science and Technology, Thuwal 23955, Saudi Arabia.

## Abstract

**Motivation:** Drug repurposing is the problem of finding new uses for known drugs, and may either involve finding a new protein target or a new indication for a known mechanism. Several computational methods for drug repurposing exist, and many of these methods rely on combinations of different sources of information, extract hand-crafted features and use a computational model to predict targets or indications for a drug. One of the distinguishing features between different drug repurposing systems is the selection of features. Recently, a set of novel machine learning methods have become available that can efficiently learn features from datasets, and these methods can be applied, among others, to text and structured data in knowledge graphs.

**Results:** We developed a novel method that combines information in literature and structured databases, and applies feature learning to generate vector space embeddings. We apply our method to the identification of drug targets and indications for known drugs based on heterogeneous information about drugs, target proteins, and diseases. We demonstrate that our method is able to combine complementary information from both structured databases and from literature, and we show that our method can compete with well-established methods for drug repurposing. Our approach is generic and can be applied to other areas in which multi-modal information is used to build predictive models.

**Availability:** https://github.com/bio-ontology-research-group/multi-drug-embedding

## 1 Introduction

The process of finding a new drug that binds a specific protein or can be used to treat a specific disease is usually time consuming and costly, taking many years and often millions of dollar (Paul *et al.*, 2010). In response, several computational approaches have been developed to identify drug targets and indications for known drugs (Chen *et al.*, 2015; Pryor and Cabreiro, 2015). Many of these approaches utilize the large volumes of data that have become available in the public domain about chemical compounds, drug and protein structures, or target sequences.

Computational drug repurposing and drug discovery methods include chemoinformatics-based methods, network-based methods, and methods based on data- or text-mining. Chemoinformatics-based methods include panel docking and ligand-based approaches (Katsila *et al.*, 2016; Meng *et al.*, 2011) which often rely on knowledge or predictions about the tertiary structure of the target protein. Network-based approaches for drug repurposing utilize different data sources, including genomic and chemical similarities and various other drugs and proteins interactions profiles or descriptors (Yamanishi *et al.*, 2008; Wang *et al.*, 2014a), integrate information related to drug mechanisms, and use machine learning techniques or graph inference methods to predict novel drug targets (Seal *et al.*, 2015; Fu *et al.*, 2016; Chen *et al.*, 2012). Additionally, omics data, in particular gene expression has been used for analyzing or inferring new drugs indications (Subramanian *et al.*, 2017). Some approaches to drug repurposing rely on data- and text-mining and are based on identification of patterns in databases or natural language text to predict novel associations between drugs and targets or drugs and diseases (Swanson, 1990; Andronis *et al.*, 2011; Frijters *et al.*, 2010; Agarwal and Searls, 2008).

The different computational approaches to drug repurposing differ both in the algorithms they employ as well as in the data sources they utilize. Finding innovative ways to use novel kinds of data or combine different types of multi-modal information in a single model has the potential to significantly improve predictive performance. For example, the PREDICT method (Gottlieb *et al.*, 2011) combined several different types of data, including similarity between proteins based on their sequence and based on their functions, as well as similarity between drugs based on their side effects or their targets. Other approaches combine disease genes associations, drugs targets, signaling pathways, and gene expression profiles to discover new therapeutic roles for known drugs (Peyvandipour *et al.*, 2018).

Many types of data that are relevant for drug repurposing can be integrated through Semantic Web technologies (Berners-Lee *et al.*, 2001), notably using the Resource Description Framework (RDF) (Manola *et al.*, 2004) and ontologies formalized in the Web Ontology Language (OWL) (Grau *et al.*, 2008). Initiatives that aim to integrate knowledge relevant for drug discovery and drug repurposing include the Open PHACTS (Williams *et al.*, 2012) and Bio2RDF (Belleau *et al.*, 2008) projects, and other relevant RDF datasets are made available by the European Bioinformatics Institute (EBI)(Jupp *et al.*, 2014).

Recently, several unsupervised machine learning methods have become available that can learn feature representations of entities represented in different types of data (LeCun *et al.*, 2015). For unstructured text, vector space models such as Word2Vec (Mikolov et al., 2013) or GLOVE (Pennington *et al.*, 2014) can learn representations of words that preserve some the words’ semantics under certain vector operations and can therefore be used to build predictive models. Similar methods have been developed for information represented as graphs (Perozzi *et al.*, 2014), knowledge graphs (Nickel *et al.*, 2016b), or formal knowledge bases (Gutiérrez-Basulto and Schockaert, 2018), and these methods have led to the development of machine learning models that can significantly outperform classic predictive methods (Alshahrani *et al.*, 2017).

Combining different types of data in a single predictive model has the potential to increase the performance of computational drug repurposing methods. Such a combination can either be done by applying different method individually on different types of data and utilize their prediction results in a new model, or by combining the different approaches within a single model.

Here, we present an approach to systematically integrate multi-modal information from knowledge graphs and literature in predictive machine learning models. Specifically, our approach utilizes structured, semantic information that can be represented in knowledge graphs and combines this information with features extracted from unstructured text. We apply the resulting combined features for drug repurposing. Using supervised machine learning, we demonstrate that our approach and the resulting model outperforms the use of individual features and leads to significant improvements when applied to computational drug repurposing.

## 2 Methods

### 2.1 Data sources

We use a knowledge graph containing information about genes/proteins, drugs, diseases, and functions generated to demonstrate the utility of knowledge graph embedding methods in life sciences (Alshahrani *et al.*, 2017). The knowledge graph is illustrated in Supplementary Figure 1. This graph consists of three ontologies, the Gene Ontology (GO) (Ashburner *et al.*, 2000), Disease Ontology (DO) (Schriml *et al.*, 2011), and the Human Phenotype Ontology (HPO) (Köhler *et al.*, 2014). It also includes three types of biological entity: diseases, genes or proteins (we do not distinguish between them in our graph), and chemicals or drugs, as well as their interactions or associations with ontology classes. The graph further includes relations between entities such as the interactions between genes/proteins obtained from STRING (Szklarczyk *et al.*, 2011) (file `protein.actions.v10.txt.gz`), chemical–protein interactions from STITCH (Kuhn *et al.*, 2012) (file `9606.actions.v4.0.tsv`), and drugs and their indications from SIDER (Kuhn *et al.*, 2015) (file `meddra_all_indications.tsv`). We built the graph using RDF and downloaded all evaluation data on 11 March 2018.

For text processing, we use the pre-annotated Medline corpus provided by the PubTator project (Wei *et al.*, 2013), downloaded on 18 Dec 2017. This corpus contains 27,599,238 abstracts together with annotations to chemicals, genes/proteins, and diseases. We use the annotations provided by PubTator for chemicals, genes/proteins, and diseases. PubTator has annotations to 17,505,118 chemicals mentions covering 129,085 distinct drugs using either CHEBI or MESH identifiers. We could map 9,545 of these to STITCH identifier using the file `9606.protein.aliases.v10.txt` provided by STITCH. PubTator further contains 81,655,248 disease mentions covering 8,143 distinct diseases in MESH. We use the DO and map these to 2,581 distinct DO classes. Furthermore, PubTator contains 17,260,141 gene/protein mentions covering 137,353 distinct genes in different species, 35,466 of which refers to human genes.

### 2.2 Generation of corpus and text normalization

We use an edge-labeled iterated random walk of fixed length and without restart to generate a corpus from the knowledge graph (Alshahrani *et al.*, 2017). For each vertex in the graph, we generate a sentence based on a short random walk. Each walk is a sequence of tokens, i.e., nodes and edges. We have two parameters for corpus generation: walk length and number of walks. Walk length is the size of each walk sequence and the number of walks is the total number of walks generated for each vertex. For all experiments, we use a walk length of 20 and perform 50 random walks for each node.

### 2.3 Learning Embeddings

We use Word2Vec (Mikolov et al., 2013) to generate embeddings for entities in our knowledge graph and for words found in text. Word2vec is a vector space model mapping words to vectors based on the co-occurrence of a word with other words within a context window across a corpus of text. In our graph, this semantics is captured by the random walks representing the co-occurrence of different entities and relations. We use the skip-gram model (Mikolov et al., 2013) in Word2Vec on the corpus generated by random walks on the knowledge graph. As parameters for both corpora, we use negative sampling using 5 words drawn from the noise distribution, a window size of 10, and an embedding size of 128.

### 2.4 Training of supervised prediction models

We evaluated the performance of each embedding method by using the embedding vectors to predict drug–target or drug–disease associations in a supervised manner. As prediction models, we use artificial neural networks, random forests, and logistic regression. For training the neural networks model, we used an architecture with a single hidden layer consisting of twice the size of the input features vector. We use a Rectified Linear Unit (ReLU) (Nair and Hinton, 2010) as an activation function for the hidden layer and a sigmoid function as the activation function for the output layer; we use cross entropy as loss function in training, and Rmsprop (Hinton *et al.*, 2012) to optimize the neural networks parameters during training. For training the neural networks, we used the Keras library in Python (Chollet *et al.*, 2015). For training the random forest classifier, we specified the number of trees to be 50, with the minimum number of training samples in leaf nodes to be one, and Gini impurity index to measure the quality of the split. For the logistic regression classifier, we used the default settings of scikit-learn in Python (Pedregosa *et al.*, 2011).

### 2.5 Multi-modal drug repurposing

We compare the performance of our method using a validation strategy in which we randomly remove a set of associations and measure how well our method can reproduce these. For each interaction type (drug–target or drug–disease associations), we remove all interactions from our knowledge graph before generating embeddings. We then use 80% of the associations to train a classifier and we test and evaluate our method on the remaining 20%. We identify the 20% randomly among all interactions. Furthermore, we randomly select an equal number of weak negative interactions, i.e., pairs of a drug and gene/protein (when predicting targets) or disease (when predicting interactions) that are not known to be associated, and we use these as negatives during training of our classifiers.

For the rank-based evaluation (ROCAUC or recall at certain ranks), we create an embedding matrix for each drug in which we fix the first part of the matrix to represent a particular drug embedding and the second part represents the gene/protein or disease embedding. We then apply the learned model on the matrix and rank the genes/proteins or diseases based on the confidence scores provided by the classifiers.

The true positive rate (TPR) and false positive rate (FPR) at each rank are used to identify the proportion of correctly and falsely predicted interactions. We quantify the performance of the predictions through the area under the receiver operating characteristic (ROC) curve (Fawcett, 2006). A ROC curve is a plot of the TPR as a function of the FPR. The TPR at a particular rank is defined as a rate of correctly predicted drug– target interactions or drug–disease associations, and the FPR is the rate of predicted interactions that are not drug–target or drug–disease interactions. As we do not have true negative drug–target interactions, we use “weak” negatives and treat all unassociated pairs of drug and gene/protein or disease as negatives. The recall at ranks ten and 100 is calculated as the ratio of predicted true positives in the top ten or 100 over the total number of positives.

## Results

### 3.1 Integrating literature and structured knowledge

Information about drugs and their targets is present in several locations and formats, including in structured databases and in scientific literature. We base our approach on an integrated dataset consisting of structured data from multiple databases and literature. We use the Resource Description Framework (RDF) (Beckett, 2004) to express and integrate structured information we consider useful for predicting drug–target and drug– indications associations. In RDF, knowledge is expressed in a graph-based format in which entities are represented by an Internationalized Resource Identifier (IRI) and relations between entities as a property that connects two nodes.

We integrate several datasets related to drug actions and diseases in a knowledge graph using RDF as representation language. Specifically, we combine information about drugs and their targets (Kuhn *et al.*, 2012) and indications (Kuhn *et al.*, 2015), gene–disease associations (Piñero *et al.*, 2017), and disease phenotypes (Hoehndorf *et al.*, 2015), as well as gene functions and interactions between gene products (Szklarczyk *et al.*, 2011). For example, we link the disease *Primary pulmonary hypertension* (`DOID:14557`) to the phenotype *Arrhythmia* (`HP:0011675`) (using a `has phenotype` relation), we link the gene *CAV1* to disease *Primary pulmonary hypertension* (`DOID:14557`) (using a `has disease association` relation), and we link the drug *Tadalafil* (`CID00110635`) to phenotype *Abdominal pain* (`HP:0002027`) (using a `has sideeffect` relation):

@prefix doid: <http://purl.obolibrary.org/obo/DOID_>.

@prefix hp: <http://purl.obolibrary.org/obo/HP_>.

@prefix b2v: <http://bio2vec.net/relation/>.

@prefix entrez: <http://www.ncbi.nlm.nih.gov/gene/>.

@prefix stitch: <http://bio2vec.net/CID>.

`doid:14557 b2v:has_disease_phenotype hp:0011675.`

`entrez:857 b2v:has_disease_association doid:14557.`

`stitch:00110635 b2v:has_sideeffect hp:0002027.`

`stitch:00110635 b2v:has_indication doid:65.`

We further add biological background knowledge expressed in ontologies, specifically the Human Phenotype Ontology (HPO) (Köhler *et al.*, 2014), Gene Ontology (GO) (Ashburner *et al.*, 2000) and Disease Ontology (DO) (Schriml *et al.*, 2011), directly to this RDF graph so that the superclasses of phenotypes can be accessed and used during machine learning. Supplementary Figure 1 shows the graph we build and the relations between the different biological entities it includes.

To learn representations of features contained in this knowledge graph, we apply a random walk algorithm over RDF and OWL (Alshahrani *et al.*, 2017) and generate a corpus consisting of iterated random walks through this graph, starting from each node. We consider each random walk as a “sentence” that expresses a chain of statements following a random path through the knowledge graph.

As next step, we integrate the information in our knowledge graph with information contained in biomedical literature. For this purpose, we normalize biomedical literature abstracts to our knowledge graph using named entity recognition and entity normalization approaches (Rebholz-Schuhmann *et al.*, 2012) that were developed for the entities in our graph. Specifically, we normalize drug, gene, and disease mentions to our graph using the literature annotations of PubMed abstracts provided by the PubTator (Wei *et al.*, 2013) database as well as mappings provided between different vocabularies of drugs and diseases (see Methods). PubTator aggregates different entity normalization approaches such as GNorm (Wei *et al.*, 2015) or DNorm (Leaman *et al.*, 2013), which can also be used directly with new text.

As a next step, we process the annotated corpus of PubMed abstracts by replacing each mention of an entity (i.e., gene, chemical compounds, or disease) that is also included in our knowledge graph with the IRI used to represent the entity in the knowledge graph. Specifically, we ensure that the mentions in text that can be normalized to our knowledge graph are “token-identical” to the entities we represent in our knowledge graph (and the corpus resulting from our random walks through the graph). This replacement ensures that our text corpus and knowledge graph overlap on the level of tokens, and we can use this overlap in our machine learning models. Figure 1 illustrates the normalization step between the text corpus and knowledge graph.

**Fig. 1.**
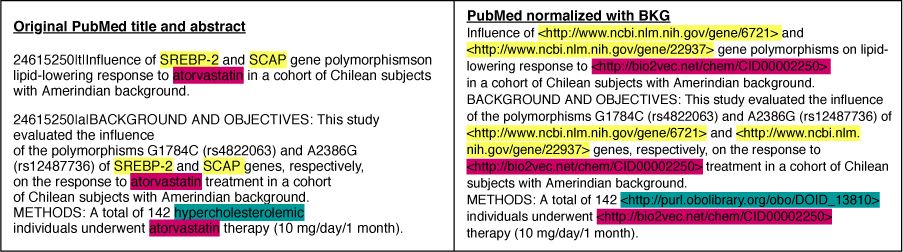
Illustration of how we normalize literature abstracts to our knowledge graph to ensure that both overlap on the level of tokens.

As an end result of these processing steps we have generated two corpora: one consisting of random walks starting from nodes in our knowledge graph, and another consisting of literature abstracts in which the mentions of entities that also appear in our graph have been replaced by the entities’ IRI in the graph. These two corpora form the foundation of our feature learning step.

### 3.2 Learning and combining features

Biological literature and the information in our knowledge graph will contain different information, and our aim is to establish a way to combine information in both data sources within a single predictive model. To achieve this goal, we first apply an unsupervised machine learning approach to generate *embeddings* for entities in our knowledge graph as well as for the words in our text corpus. An embedding encodes within a vector space the context in which an entity appears; since we represent information about entities in the knowledge graph through sentences generated from random walks, we apply the Word2Vec skip-gram model (Mikolov et al., 2013) to generate embeddings for all terms that occur within the two corpora we generated.

We use two different approaches to combine the embeddings from the knowledge graph and text corpus: first, we generate embeddings independently on both corpora, and concatenate the resulting embedding vectors; and second, we concatenate the two corpora and generate embeddings from the combined corpus. As a result, we obtain embeddings for drugs, genes, diseases, and for all other entities from the knowledge graph; we further obtain embeddings for all words that are used in our literature corpus, in particular for the entities which are mentioned in literature and which we normalized to our knowledge graph. However, not all entities in our knowledge graph also have a representation in literature, and not all entities (drugs, diseases, and genes) mentioned in literature are included in our knowledge graph. Supplementary Figure 2 shows the overlap between the two datasets. The embedding vectors generated for the entities from the two different corpora (either individually or jointly) form our entities’ feature representations, and the features either represent information from structured databases, from literature, or both. Supplementary Figure 3 and 4 show a visualization of the embeddings (from the knowledge graph, literature, and combined) using t-SNE (van der Maaten and Hinton, 2008), and coloring disease embeddings based on their top-level DO class, and drug embeddings based on their top-level class in the Anatomical Therapeutic Chemical (ATC) Classification System.

### 3.3 Evaluation on prediction of drug targets and indications

Our method combines heterogeneous information from text and a knowledge graph in vector space embeddings, and we evaluate the performance of our method by predicting drug targets and drug indications. For this purpose, we use four different evaluation methods: first, we use the embeddings generated from the knowledge graph alone; second, we use the embeddings generated from our literature corpus alone; third, we concatenate the embeddings from the knowledge graph and text corpus; and finally, we combine the text corpus and knowledge graph corpus and learn joint embeddings.

We use as evaluation sets the drug targets from the STITCH database (Kuhn *et al.*, 2012) and drug indications from the SIDER database (Kuhn *et al.*, 2015). Furthermore, to clearly distinguish and evaluate the contributions of the different data sources, we initially limit our evaluation set to the drugs, targets, and diseases that have a representation both in our knowledge graph and in our literature corpus. For predicting drug targets, we use as evaluation set 820 drugs and rank 17,380 genes that are both in our knowledge graph and found in the literature corpus. For predicting drug indications, we use 754 drugs with one or more known indications and rank 2,552 diseases (overlapping between literature and our knowledge graph) for each of the drugs to determine for which disease it may be indicated (see Supplementary Tables 1 and 2 for details).

To predict associations between drugs and their targets or drugs and the diseases they may treat, we use supervised machine learning and train a model based on 80% randomly chosen drug–target or drug–disease associations and test whether the model is able to predict the remaining 20%. We use three different machine learning approaches for the model construction: logistic regression, a random forest classifier, and artificial neural networks. Before training the model, we remove all `has-target` (when predicting drug targets) or `has-indication` (when predicting drug indications) edges in the graph before generating the corpus for predicting drug targets and indications, respectively. Each model has as input two embedding vectors that represent a drug and another embedding vector representing either a gene/protein (when predicting targets) or disease (when predicting indications). The models are trained as binary classifiers and output whether the drug targets the gene/protein or treats the disease. Figure 2 provides an overview over our overall workflow.

**Fig. 2.**
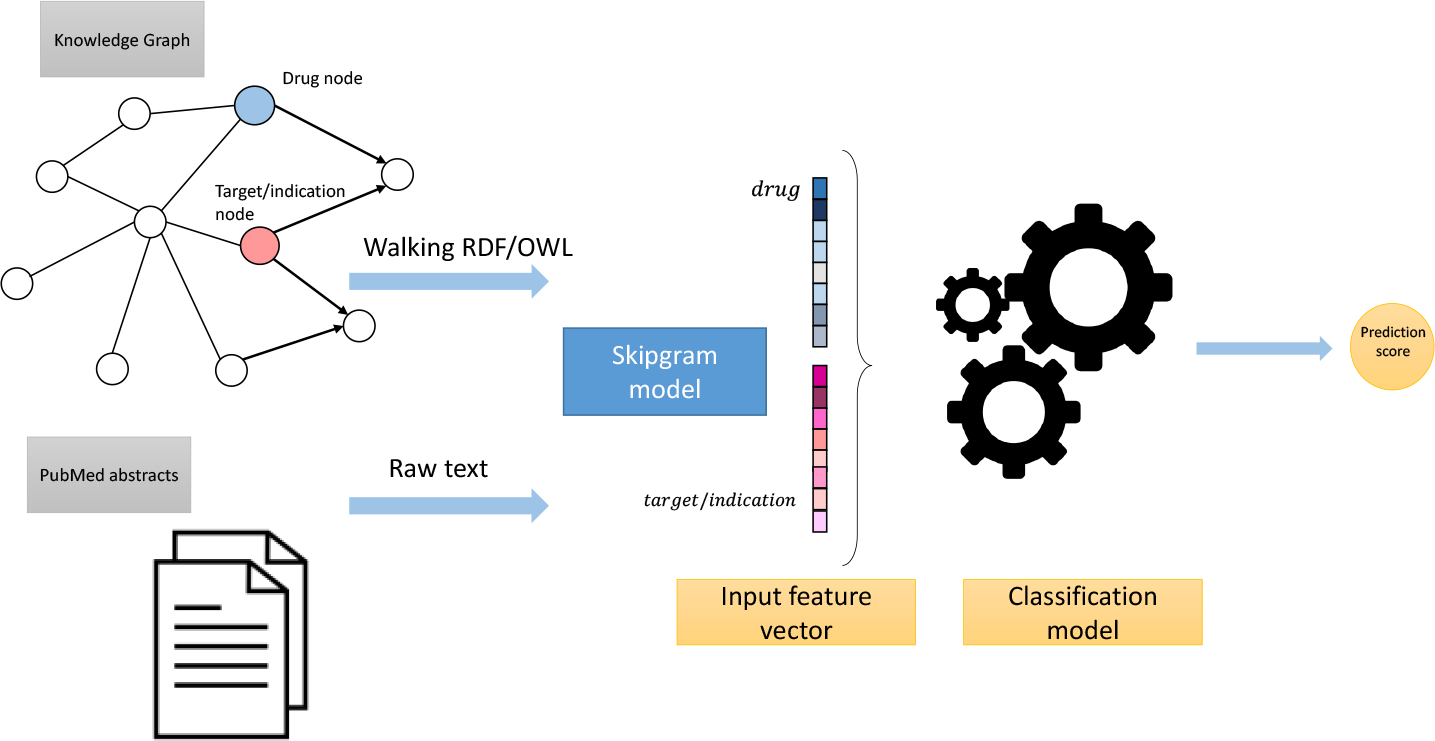
High-level overview over our workflow.

We evaluate the performance of each model on the 20% of associations we withhold from training. All three of our classification models can provide confidence values for a prediction, and we rank predicted associations based on their confidence value. We then calculate the area under the receiver operating characteristic (ROC) curve (ROCAUC) (Fawcett, 2006) as well as the number of correct associations we retrieve within the first ten and first 100 ranks. Table 1 summarizes our results for predicting associations between drugs and targets, and Table 2 summarizes the results for predicting indications.

**Table 1.**
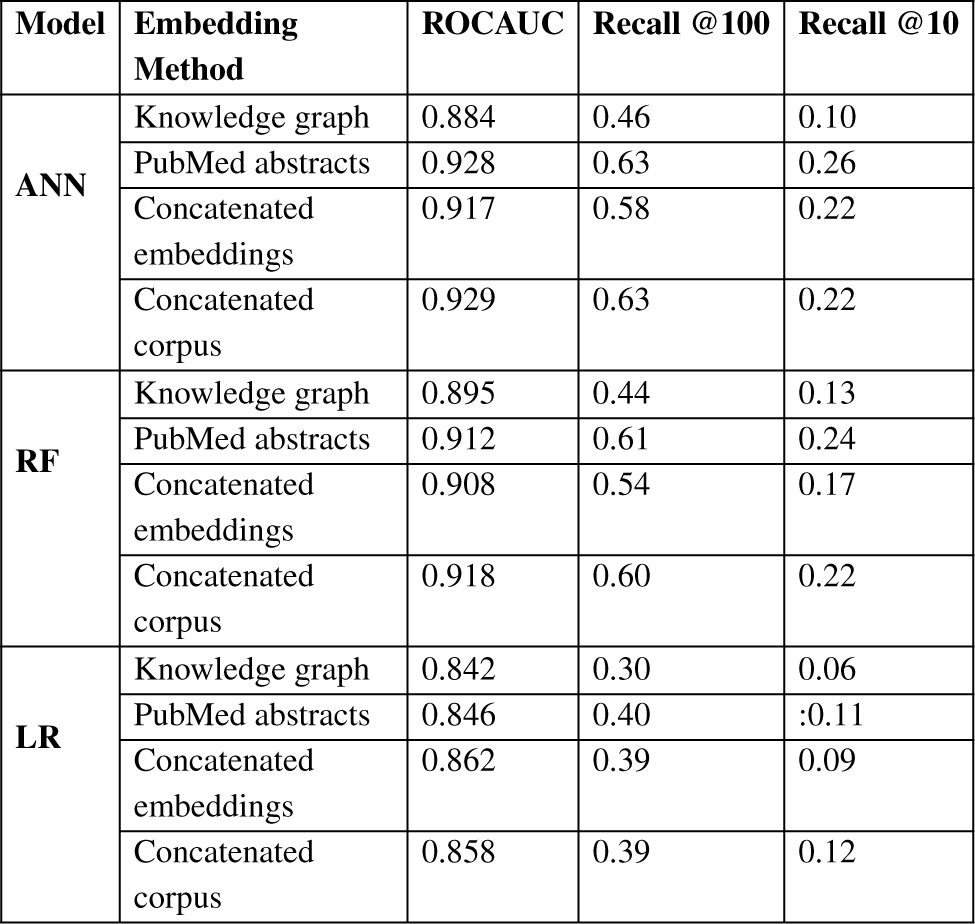
Performance results for predicting drug-target associations, based on our four embeddings approaches and using three classification models (Artificial Neural Networks (ANN), Random Forest (RF) and Logistic regression (LR)).

**Table 2.**
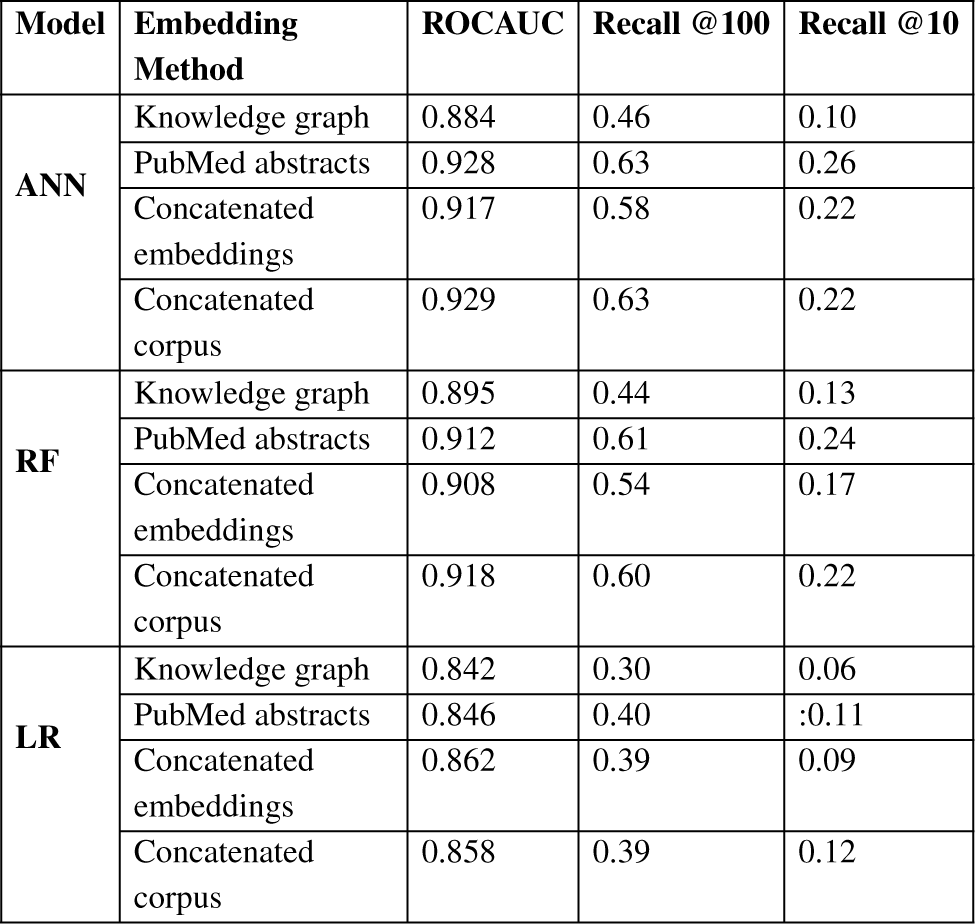
Performance results for prediction of drug indications, based on our four embeddings approaches and using three classification models (Artificial Neural Networks (ANN), Random Forest (RF) and Logistic regression (LR)).

We find that both our artificial neural network and the random forest classifier are able to accurately predict both drug targets and drug indications, while the logistic regression classifier results in relatively worse performance. An obvious explanation is that logistic regression mainly assigns weights to individual features and does not have the ability to *compare* or *match* elements of the two input embedding vectors, while both the random forest classifier and an artificial neural network are able to provide a classification based on comparing elements of the input embedding vectors. Furthermore, we find that, in general, using embeddings generated from literature results in higher predictive performance across all classifiers compared to embeddings generated from the knowledge graph alone. Combining the embeddings, either through concatenation of the embeddings or through concatenation of the two corpora sometimes but not always improves or changes the predictive performance.

While our results indicate that both literature-derived and knowledge graph embeddings can be used to predict interactions, the main contribution of our multi-modal approach is the increased coverage through combining database content and literature (see Supplementary Figure 2). To demonstrate this application of our method, we extend our evaluation set to contain all the drugs, genes, and diseases that are found in either our knowledge graph, the literature abstract, or the union of the entities in the knowledge graph and literature trained on the combined corpus. Figure 3 shows the ROC curves and the ROCAUC for predicting drug targets and drug indications using our neural network classifier, based on a combination of the literature corpus and the random walk corpus.

**Fig. 3.**
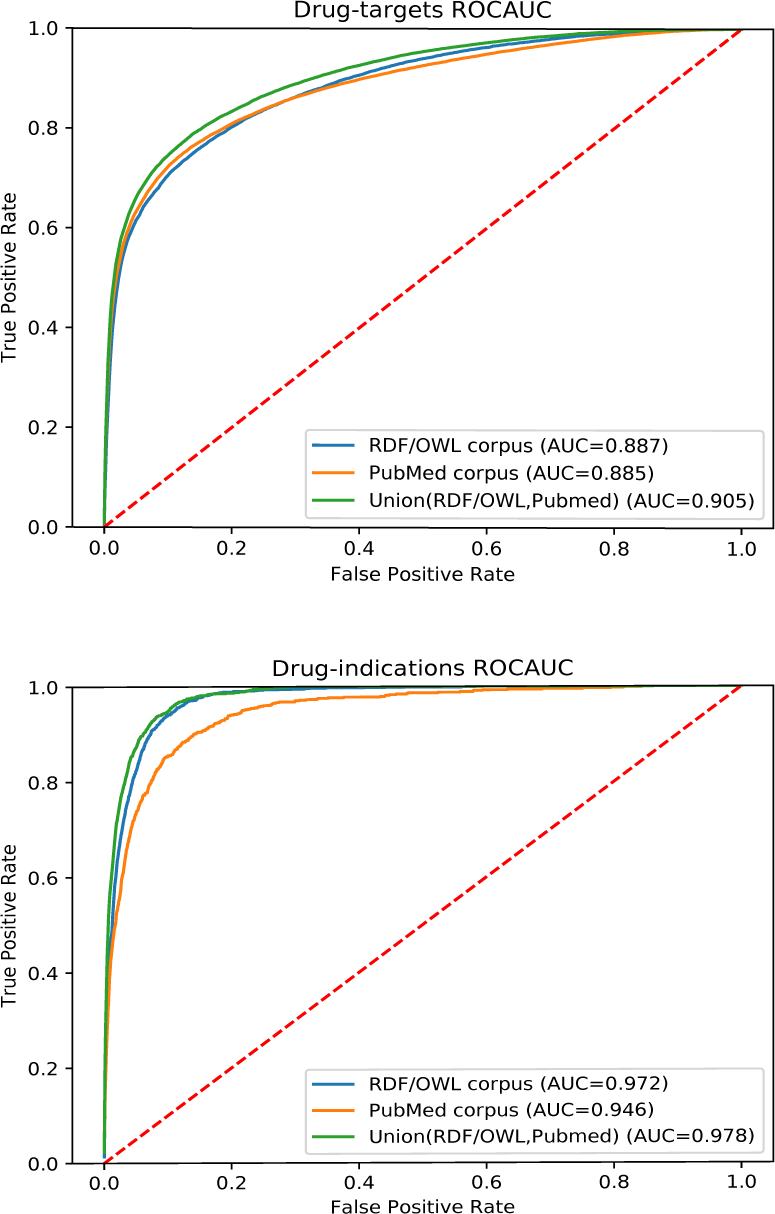
ROC curve of our neural network for predicting drug targets in the union of associations present in the knowledge graph and PubMed abstracts (top); ROC curve of our neural network for predicting drug indications found in the union of knowledge graph and PubMed abstracts.

Our knowledge graph contains a very large number of chemicals, many of which are not drug-like, and while the performance in predicting drug targets is somewhat higher when using the knowledge graph embeddings, the overall performance is still dominated by the literature-derived embedding vectors. However, when predicting indications for known drugs, both our graph and literature overlap more substantially while nevertheless containing complementary information. We observe a significant improvement in predicting drug indications when combining the information from literature and the knowledge graph. We make all predictions of drug–target associations as well as the predictions for drug indications freely available at https://github.com/bio-ontology-research-group/multi-drug-embedding.

## 4 Discussion

There are many scenarios in biological and biomedical research in which predictive models need to be built that can utilize information that is represented in different formats. Our key contribution is a method to integrate data represented in structured databases, in particular knowledge graphs represented in RDF and OWL, and integrate this information with information in literature. While we primarily focus on the prediction of drug-target interactions and drug indications based on information in text and databases, our approach is generic and can serve as a paradigm for learning from multi-modal, heterogeneous data in biology and biomedicine.

Our method uses feature learning to project different types of data into a vector space, and combine data of different modes either within a single vector space (when mapping data of different modes to the same space, or to vector spaces of identical dimensions) or we combine the vector spaces themselves. We rely on the recent success of deep learning methods (Ravì *et al.*, 2017; Angermueller *et al.*, 2016) which improved our ability to learn relevant features from a data set and project them into a vector space. In particular, our approach relies on natural language models, in particular Word2Vec (Mikolov et al., 2013), and recent approaches to project information in knowledge graphs into vector spaces (Nickel *et al.*, 2016b; Alshahrani *et al.*, 2017; Smaili *et al.*, 2018). These approaches are now increasingly applied in biological and biomedical research (Alshahrani and Hoehndorf, 2018) yet often restricted to single types of representation (such as images, genomic sequences, text, or knowledge graphs).

Our approach naturally builds on the significant efforts that have been invested in the development of named entity recognition and normalization methods for many different biological entities (Rebholz-Schuhmann *et al.*, 2012) as well as the effort to formally represent and integrate biological data using Semantic Web technologies (Jupp *et al.*, 2014; Callahan *et al.*, 2013). Several biological data providers now provide their data natively using RDF (Jupp *et al.*, 2014; UniProt Consortium, 2018). Furthermore, many methods and tools have been developed to normalize mentions of biological entities in text to biological databases, for example for mentions of genes and proteins, (Leaman and Gonzalez, 2008; Wei *et al.*, 2015), chemicals (Leaman *et al.*, 2015) as well as diseases (Leaman *et al.*, 2013), and repositories have been developed to aggregate and integrate the annotations to literature abstracts or fulltext articles (Wei *et al.*, 2013; Kim and Wang, 2012). While these methods, tools, and repositories are not commonly designed to normalize mentions of biological entities to a knowledge graph, we demonstrate here how a normalization of text to a knowledge graph can be achieved, and subsequently use the combined information in our multi-modal machine learning approach. Consequently, our method has the potential to increase the value of freely available Linked Data resources and connect them directly to the methods and tools developed for natural language processing and text mining in biology and biomedicine.

In the future, it would be beneficial to develop better entity normalization methods that can directly normalize entity mentions in text to a knowledge graph. We also intend to evaluate the success of our approach on full-text articles so that more information, in particular regarding methods and experimental protocols, can be utilized by our approach. Methodologically, we also intend to apply other knowledge graph embedding methods, in particular translational embeddings (Bordes et al., 2013; Nickel *et al.*, 2016a; Dai and Yeung, 2006), that have previously been combined successfully with textual information (Wang *et al.*, 2014b), and evaluate their performance for prediction of biological relations.

